# Stress contingent changes in Hog1 pathway architecture and regulation in *Candida albicans*

**DOI:** 10.1101/2024.06.05.597528

**Authors:** Alison M. Day, Min Cao, Alessandra da Silva Dantas, Carmen Herrero-de-Dios, Alistair J. P. Brown, Janet Quinn

## Abstract

The Hog1 stress-activated protein kinase (SAPK) is a key mediator of stress resistance and virulence in *Candida albicans*. Hog1 activation via phosphorylation of the canonical TGY motif is mediated by the Pbs2 MAPKK, which itself is activated by the Ssk2 MAPKKK. Although this three-tiered SAPK signalling module is well characterised, it is unclear how Hog1 activation is regulated in response to different stresses. Functioning upstream of the Ssk2 MAPKKK is a two-component related signal transduction system comprising three sensor histidine kinases, a phosphotransfer protein Ypd1, and a response regulator Ssk1. Here, we report that Ssk1 is a master regulator of the Hog1 SAPK that promotes stress resistance and Hog1 phosphorylation in response to diverse stresses, except high osmotic stress. Notably, we find Ssk1 regulates Hog1 in a two-component independent manner by functioning as a scaffolding protein to promote interactions between the Ssk2 and Pbs2 kinases. We propose this scaffolding function is important to maintain a basal level of Hog1 phosphorylation which is necessary for oxidative stress, but not osmotic stress, mediated Hog1 activation. We find that osmotic stress triggers robust Pbs2 phosphorylation which drives its dissociation from Ssk2. In contrast, Pbs2 is not robustly phosphorylated following oxidative stress and the Ssk1-mediated Ssk2-Pbs2 interaction remains intact. Instead, oxidative stress-stimulated increases in phosphorylated Hog1 is dependent on the inhibition of protein tyrosine phosphatases that negatively regulate Hog1 coupled with the Ssk1-mediated promotion of basal Hog1 activity. Furthermore, we find that inhibition of protein tyrosine phosphatases is linked to the hydrogen peroxide induced oxidation of these negative regulators in a mechanism that is dependent thioredoxin. Taken together these data reveal stress contingent changes in Hog1 pathway architecture and regulation and uncover a novel mode of action of the Ssk1 response regulator in SAPK regulation.

**Author summary:** As a core stress regulator, the Hog1 stress-activated protein kinase (SAPK), is a key virulence determinant in many fungal pathogens. Despite this, little is known regarding the mechanisms by which different stresses trigger the phosphorylation and activation of Hog1. Here we present three novel findings regarding Hog1 regulation in the human fungal pathogen *C. albicans*. Firstly, we find that the response regulator protein, Ssk1, is a master regulator of Hog1 that forms a scaffold for the upstream Hog1-activating kinases, Ssk2 and Pbs2. Secondly, this scaffolding role maintains a basal level of Hog1 phosphorylation, which is important for responses to stresses, such as oxidative stress, that do not stimulate activation of the upstream Ssk2 and Pbs2 kinases. Instead, oxidative stress induced Hog1 phosphorylation is mediated through the oxidation and inactivation of protein tyrosine phosphatases that negatively regulate Hog1. Finally, we show that high osmotic stress induces the robust phosphorylation and activation of the upstream kinase Pbs2, which drives its dissociation from the Ssk1-mediated scaffold. These new insights into the regulation of the *C. albicans* Hog1 SAPK pathway offer new strategies to therapeutically target this core virulence determinant.

## Introduction

*Candida albicans* is categorized as a critical fungal priority pathogen by the World Health Organization (1). As a common commensal of the human gut mycobiome, *C. albicans* is an opportunistic pathogen that is poised to cause serious invasive infections in immunocompromised hosts (2). Treatment options are limited with only four classes of drugs licensed to treat systemic fungal infections (3), and emerging resistance to these drugs is an increasing problem (4). This, coupled with the fact that candidemia is associated with a crude mortality rate of between 25-40% (5, 6), underscores the need to develop new strategies to prevent life-threatening *C. albicans* infections.

The ability of *C. albicans* to sense and adapt to changing and often harsh environments during disease progression is a key virulence trait (7, 8). One of the most hostile host environments encountered during infection is during phagocytic attack by innate immune cells where the fungus is subjected to a cocktail of antimicrobial agents including toxic reactive oxygen and nitrogen species, cationic and toxic metal influxes, and antimicrobial peptides and digestive enzymes, all within a nutrient poor and highly acidic environment (9).

*C. albicans* also encounters different environmental stresses depending on the host niche occupied (8). For example, changes in pH are encountered upon moving from the intestine (pH range 5-6.5) to the bloodstream (pH 7.4) (10), high salinity levels are encountered in the kidneys (11), and nutritional immunity strategies either restrict access to, or expose the fungus to toxic levels of divalent cations (12).

The Hog1 stress-activated protein kinase (SAPK) is a core stress-signalling protein that senses and responds to many physiologically relevant stress conditions (reviewed in (13)). Hog1 promotes *C. albicans* resistance to numerous stresses including oxidative and nitrosative stresses, osmotic and cationic stresses, antimicrobial peptides, heavy metals, various antimicrobial drugs, and ER stress (13). In addition to stress resistance, Hog1 is implicated in several morphological and phenotypic transitions in *C. albicans* (14, 15), and is also linked to cell size control, respiratory metabolism, and induction of macrophage pyroptosis ((16–18). Consistent with the multi-faceted roles of this central signalling hub, Hog1 is essential for *C. albicans* virulence in models of systemic infection (14, 19), colonisation in models of gut commensalism (20), and for survival following phagocytosis by macrophages and neutrophils (21, 22).

Hog1-related SAPKs are found in all eukaryotic cells and are amongst the most evolutionarily conserved stress-signalling proteins (23). Activation of the SAPK is mediated through phosphorylation of conserved threonine and tyrosine residues within a TGY motif located in the activation loop of the kinase (24). In *C. albicans,* stress-induced Hog1 phosphorylation occurs in response to many of the stresses outlined above. This triggers the nuclear accumulation of Hog1 and the subsequent phosphorylation of both nuclear and cytoplasmic substrates drives appropriate cellular responses (13). As seen in all SAPK modules, the Hog1 pathway in *C. albicans* comprises three tiers of sequentially acting protein kinases; the MAPKKK Ssk2, which phosphorylates and activates the MAPKK Pbs2, which in turn phosphorylates and activates Hog1 (19, 25, 26). Accordingly, stress induced phosphorylation of Hog1 is abolished in *ssk2Δ* and *pbs2Δ* cells which in turn display overlapping stress-sensitive phenotypes to those exhibited by *hog1Δ* cells (25, 26). Hog1 phosphorylation is also negatively regulated by the serine/threonine phosphatases Ptc1 and Ptc2 (18) and the tyrosine phosphatases Ptp2 and Ptp3 (27). Thus, the level of Hog1 phosphorylation is dynamically regulated by the opposing action of upstream kinases and downstream phosphatases.

Although Hog1 is essential for virulence, the presence of structural and functional homologues in human cells, such as p38, has reduced the attractiveness of Hog1 as an antifungal target. Nevertheless, there has been interest in identifying fungal-specific regulators of SAPKs as potential drug targets (28). Such fungal-specific mechanisms include two-component related signalling pathways that regulate the activity of MAPKKKs within the Hog1 pathway. In *C. albicans* the two-component pathway comprises of three sensor histidine protein kinases (Sln1, Chk1 and Nik1), which relay phosphate to an intermediary phosphotransfer protein (Ypd1), which in turn transfers phosphate to an aspartate residue within the response regulator Ssk1 (reviewed in (29)). Deletion of the Ypd1 phosphorelay protein in *C. albicans* results in the constitutive phosphorylation and hyperactivation of the Hog1 kinase in a mechanism that is dependent on the downstream response regulator Ssk1

(30). This is consistent with work in *S. cerevisiae* showing that unphosphorylated Ssk1 is a potent activator of the Ssk2/Ssk22 MAPKKKs (31). In *C. albicans*, deletion of Ssk1 was found to impact Hog1 activation in response to oxidative stress, but not osmotic stress (32). This is intriguing as none of the histidine kinases appear to relay oxidative stress signals to Hog1 (33) suggesting that Ssk1 may also regulate Hog1 in a mechanism that is independent of two-component signalling. Indeed, oxidative stress-induced activation of Hog1 was observed in cells expressing a mutated version of Ssk1 that is no longer responsive to its upstream phosphorelay module (34).

Ssk1 is essential for *C. albicans* virulence (35), and as a fungal-specific regulator of Hog1 is a potential target for the development of antifungals. Here we provide new insight into two-component independent roles of Ssk1 in Hog1 regulation and additionally provide mechanistic insight into stress contingent changes in SAPK pathway architecture and regulation in *C. albicans*.

## Materials and Methods

### Strains and growth media

All the *C. albicans* strains used in this study are listed in Table 1. Cells were grown at 30°C in YPD rich medium (36).

**Table 1.** Strains used in this study.

| Strain | Description | Genotype | Source |
| --- | --- | --- | --- |
| RM1000 | parental | *ura3::λ imm434/ura3::λimm434, his1::hisG/his1::hisG* | (64) |
| JC21 | WT | RM1000 + Clp20 | (43) |
| JC50 | *hog1Δ* | RM1000 *hog1::loxP-ura3-loxP, hog1::loxP-HIS1-loxP* + Clp20 | (43) |
| JC112 | Pbs2-HM | RM1000 *pbs2::loxP-HIS1-loxP/PBS2-HM:URA3* | (19) |
| JC1188 | *ssk1Δ* | RM1000 *ssk1::loxP-ura3-loxP, ssk1::loxP-HIS1-loxP* + Clp10 | This study |
| JC1196 | *ssk1Δ+SSK1* | RM1000 *ssk1::loxP-ura3-loxP/ssk1::loxP-HIS1-loxP* + Clp10-SSK1 | This study |
| JC2001 | *ypd1Δ* | RM1000 *ypd1::HIS1/ypd1::hisG* + Clp20 | (30) |
| JC2119 | Pbs2<sup>AA</sup>-HM Ssk2-HA | RM1000 *pbs2::loxP-HIS1-loxP/PBS2<sup>AA</sup>-HM:URA3, SSK2-HA:ARG4* | This study |
| JC2121 | Pbs2-HM Ssk2-HA | RM1000 *pbs2::loxP-HIS1-loxP/PBS2-HM:URA3, SSK2-HA:ARG4* | This study |
| SN148 | parental | *ura3::λ imm434/ura3::λimm434, his1::hisG/his1::hisG arg4::hisG/arg4::hisG* | (39) |
| JC747 | WT | SN148 + Clp30 | (45) |
| JC1519 | *ssk1Δ* | SN148 *ssk1::HIS1/ssk1::ARG4* + Clp10 | This study |
| JC1554 | *ssk1Δtrx1Δ* | SN148 *ssk1::HIS1/ssk1::ARG4, trx1::hisG/trx1::hisG* + Clp10 | This study |
| JC1555 | *ssk1Δtrx1Δ+SSK1* | SN148 *ssk1::HIS1/ssk1::ARG4, trx1::hisG/trx1::hisG* + Clp10-SSK1 | This study |
| JC2012 | Pbs2-HM Ssk2-HA *ssk1Δ* | SN148 *PBS2-HM:URA3, SSK2-HA:ARG4 ssk1::hisG/ssk1::hisG* | This study |
| JC2017 | Pbs2-HM Ssk2-HA | SN148 *PBS2-HM:URA3, SSK2-HA:ARG4* | This study |
| JC2231 | Ptp3-HA | SN148 *PTP3-HA:HIS1* | This study |
| JC2626 | Ptp3-HA *trx1Δ* | SN148 *PTP3-HA:HIS1 trx1::hisG/trx1::hisG* | This study |
| SN250 | WT | *hisΔ1/his1Δ, leu2Δ::C.dubliniensis HIS1/leu2Δ::C.maltosa LEU2, arg4Δ/arg4Δ, URA3/ura3Δ::imm<sup>434</sup>, IRO1/iro1Δ::imm<sup>434</sup>* | (39) |
| JC2291 | *ptp2Δptp3Δ* | SN250 *ptp3Δ::C.dubliniensis HIS1/ptp3Δ::C.maltosa LEU2 ptp2Δ::URA3/ptp2Δ::ARG4* | (27) |

### Strain construction Deletion of *SSK1*

An *SSK1* deletion mutant in RM1000 cells (JC784 (37)) was passed over 5-FOA to generate the *ura3* strain JC832. To construct re-integrant control strains the *SSK1* gene plus 1000 bp of the promoter region and 200 bp of the terminator region were amplified by PCR, using the oligonucleotide primers SSK1PromF and SSK1TermR, and ligated into the BamHI site of CIp10 (38) to create Clp10-SSK1.

The CIp10 vector and CIp10-SSK1 plasmid were digested with StuI and integrated at the *RPS10* locus in JC832 cells to generate strains JC1188 and JC1196 respectively.

A second *SSK1* deletion strain was constructed in SN148 cells (39), to allow subsequent epitope tagging of *PBS2* and *SSK2*. A *SSK1* disruption cassette containing the *HIS1* gene was generated by PCR using the oligonucleotide primers SSK1delF and SSK1delR and the plasmid template pLHL2 (40). The cassette was transformed into *C. albicans* to disrupt one allele of *SSK1* (JC1367). The second copy of *SSK1* was deleted using an *ssk1::hisG-URA3-hisG* ura-blaster cassette which was generated by amplifying the 5’ and 3’ regions flanking the *SSK1* open reading frame using oligonucleotide primer pairs 5’Ssk1BamF / 5’Ssk1BamR and 3’Ssk1BamF / 3’Ssk1BamR respectively and sequentially ligating into p5921 (41). The *ssk1::hisG-URA3-hisG* disruption cassette was released from the plasmid by digestion with restriction enzymes KpnI and HindIII and transformed into JC1367 cells to generate strain JC1980. This strain was then passed over 5-FOA to recycle the *URA3* marker and generate strain JC1981.

To generate a *ssk1Δtrx1Δ* double deletion strain, *SSK1* disruption cassettes containing the *HIS1* or *ARG4* were generated by PCR using the oligonucleotide primers SSK1delF and SSK1delR and the plasmid template pLHL2 or pLAL2 (40). These were sequentially transformed into SN148 cells to delete both copies of *SSK2* and generate strain JC1402. *TRX1* was subsequently deleted following two rounds of transformation of JC1402 cells with a *trx1::hisG-URA3-hisG* ura-blaster cassette (41). Following passage over 5-FOA to recycle the *URA3* marker, the CIp10 vector and CIp10-SSK1 plasmid were digested with StuI and integrated at the *RPS10* locus in *ssk1Δtrx1Δ* cells to generate strains JC1554 and JC1555 respectively. Oligonucleotides used in this study are available on request.

### Tagging of Pbs2, Ssk2 and Ptp3

*PBS2*, expressed from its native genomic locus, was epitope tagged with a sequence encoding 6-His2-Myc. The plasmid CIp-C-PBS2HM (26) was linearised with SgrAI to target integration at the *PBS2* locus in WT (SN148) and *ssk1Δ* (JC1981) cells to generate strains JC1803 and JC2008. To tag *SSK2* and *PTP3* at their native chromosomal loci with a triple HA epitope tag, the PCR-based approach described by Lavoie *et al*. (42) was employed. To tag *SSK2*, a tagging cassette consisting of the HA-tag and the *ARG4* gene flanked by 100 base pairs upstream and downstream of the STOP codon of *SSK2* was generated by PCR using primers SSK2HAtagF and SSK2HAtagR and pFA-HA-ARG4 as template (42). The cassette was integrated into the *SSK2* locus of JC1803 and JC2008 to generate wild type (JC2017) and *ssk1Δ* strains (JC2012) expressing both Pbs2HM and Ssk2HA. The Ssk2-3HA tagging cassette was also integrated into the *SSK2* locus of Pbs2HM (JC112) and Pbs2AA-HM (JC126) expressing cells (19) to generate strains JC2121 and JC2119 respectively. To tag *PTP3*, a tagging cassette consisting of the HA-tag and the *HIS3* gene flanked by 100 base pairs upstream and downstream of the *PTP3* STOP codon was generated by PCR using primers PTP3HAtagF and PTP3HAtagR and pFA-HA-HIS3 as template (42). The tagging cassette was integrated into the *PTP3* locus of SN148 and JC1561 cells to generate wild type (JC2231) and *trx1Δ* strains (JC2626) expressing Ptp3HA.

### Stress sensitivity tests

Cells were grown at 30°C to mid-exponential phase and were then diluted in YPD medium. 1×10^3^ cells were spotted onto YPD agar containing the stress agent indicated. Plates were incubated at 30°C for 24h. All experiments were repeated a minimum of 3 times.

### Hog1 phosphorylation assays

Cells were grown to mid-exponential phase and harvested before and after exposure to the stress agent indicated. Protein extracts were prepared as described previously (43) and 50 µg total cell protein was resolved on a 10% SDS-PAGE gel. Phosphorylated Hog1 was detected by Western blot analysis using an anti-phospho-p38 antibody (Cell signalling Technology) as described previously (43). Blots were stripped and reprobed with an anti-Hog1 antibody (y-215, Santa Cruz Biotechnology) to determine total Hog1 levels. All experiments were repeated a minimum of 3 times.

### Microscopy

Differential Interference Contrast images were captured using a Zeiss Axioscope microscope as described previously (43).

### Co-immunoprecipitation Assays

Wild type (JC2017) and *ssk1Δ* (JC2012) cells expressing chromosomally tagged Ssk2-HA and Pbs2-6His-myc were grown to mid-exponential phase and samples collected before and after exposure to the stress agent indicated. Protein extracts were prepared as described previously (26). Pbs2-6His-myc was immunoprecipitated from 1000 µg of whole extracts following incubation with 40 µl of anti-c-myc agarose (Santa Cruz Biotechnology) for 2 h at 4°C. The agarose beads were washed twice with lysis buffer (50mM Tris-HCl pH 7.5, 150mM NaCl, 20mM imidazole, 0.1% NP-40, 50mM NaF, 1mM phenylmethylsulfonyl fluoride, 0.07 trypsin inhibition units/ml aprotinin, 10µg/ml leupeptin and 10µg/ml pepstatin) and once with high salt lysis buffer (50mM Tris-HCl pH 7.5, 500mM NaCl, 20mM imidazole, 0.1% NP-40, 50mM NaF, 1mM phenylmethylsulfonyl fluoride, 0.07 trypsin inhibition units/ml aprotinin, 10µg/ml leupeptin and 10µg/ml pepstatin) and resuspended in SDS-loading buffer. Proteins were resolved on a 10% SDS-PAGE gel and Western blotting was performed using an anti-myc antibody (Merck, UK). Co-immunoprecipitation of Ssk2-HA was detected using an anti-HA antibody (Merck, UK). All experiments were repeated a minimum of 3 times.

### Oxidation of Ptp3

Exponentially growing *C. albicans* cells were added to 20% (w/v) trichloroacetic acid (TCA), harvested by centrifugation and snap frozen in liquid nitrogen. Pelleted cells were suspended in 10% (w/v) TCA and disrupted using ice cold glass beads and a Mini Beadbeater-16 (Biospec products). Following cell lysis, TCA-precipitated proteins were acetone-washed before resuspension in TES buffer (100 mM Tris-HCL pH 8.0, 1 mM EDTA, 1% (w/v) SDS) containing 10 mM N-Ethylmaleimide (NEM). Equal amounts of denatured protein added to reducing or non-reducing (-2-mercaptoethanol) sample buffer, separated by SDS-PAGE on 8% acrylamide gels and transferred onto nitrocellulose membrane (GE healthcare). Membranes were stained with Revert™ 700 total protein stain (LI-COR) for normalization of protein loading following which western blotting was performed using an anti-HA antibody (H9658 Merck, UK). HA tagged protein bands were visualised using the LI-COR Odyssey Clx system with LI-COR secondary antibody (IRDye 800 CW anti-mouse) antibody. All experiments were repeated a minimum of 3 times.

## Results

### Ssk1 is a master regulator of the Hog1 SAPK pathway

To investigate the importance of the Ssk1 response regulator in relaying stress signals to the Hog1 SAPK in *C. albicans,* the stress sensitive phenotypes of congenic *ssk1Δ* and *hog1Δ* mutant strains were compared (Fig. 1A). Consistent with previous studies (43), cells lacking the Hog1 SAPK displayed pleiotropic stress sensitivities to a range of oxidative stress-inducing agents, antifungal drugs, heavy metals, the cell membrane damaging agent SDS, and cationic and osmotic stresses. Notably, *ssk1Δ* cells displayed overlapping stress-sensitive phenotypes to those exhibited by *hog1Δ* cells. Interesting exceptions were the osmotic stress agents, NaCl and sorbitol, in which cells lacking Ssk1, unlike *hog1Δ* cells, displayed wild-type levels of resistance. Such findings are consistent with previous work that also failed to detect any sensitivity of *ssk1Δ* cells to NaCl (34).

**Figure 1.**
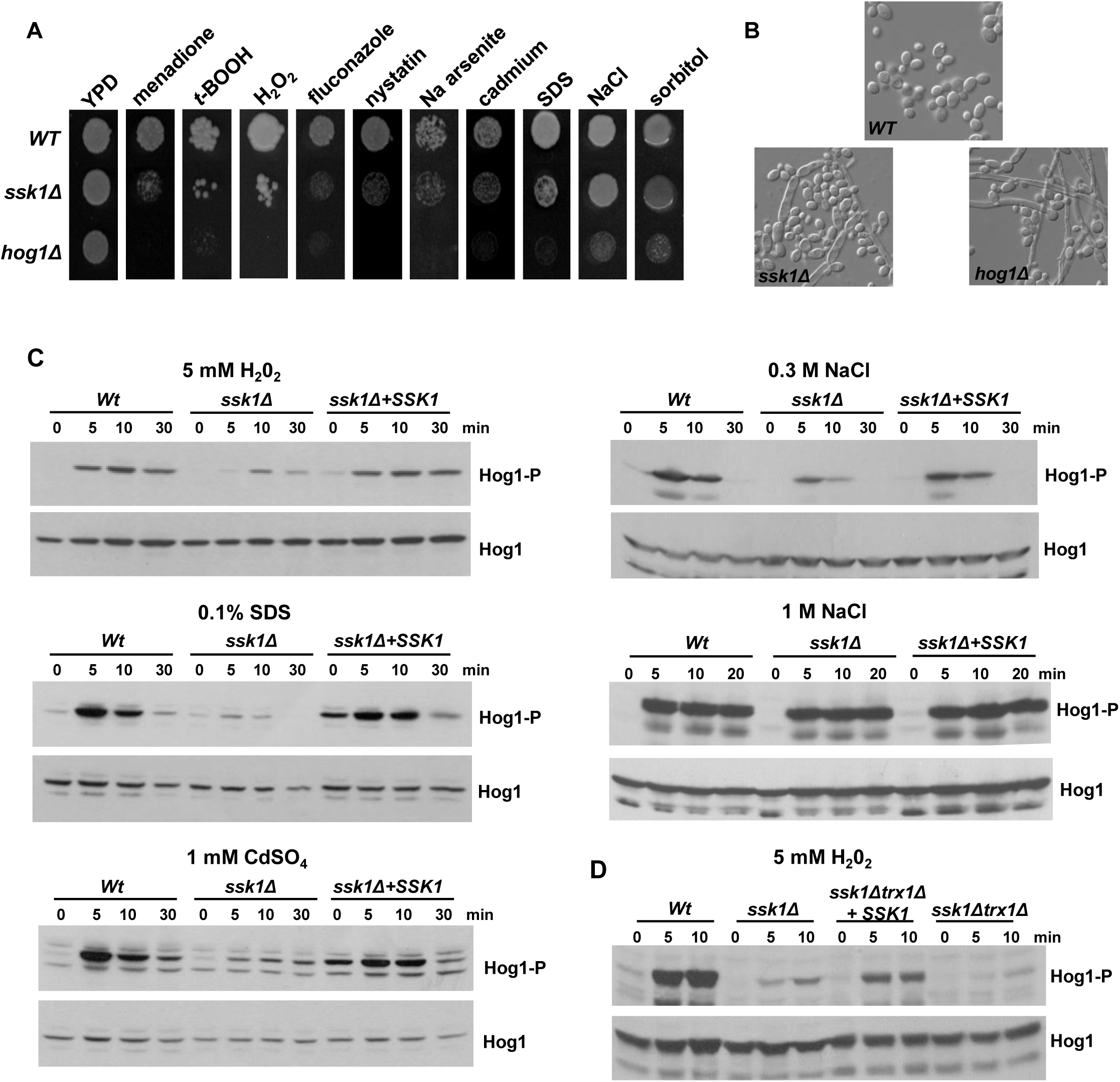
Ssk1 is a master regulator of the Hog1 SAPK pathway. (A) Approximately 1×10^3^ exponentially growing Wt (JC21), *hog1Δ* (JC50) and *ssk1Δ* (JC1188) cells were spotted onto YPD plates containing 300µM menadione, 2mM tBOOH, 3mM H_2_O_2_, 5µg/ml Fluconazole, 0.5µg/ml Nystatin, 2mM sodium arsenite, 0.4mM cadmium sulphate, 0.02% SDS, 1M NaCl or 1M sorbitol. Plates were incubated at 30°C for 48h. (B) Morphology of Wt, *hog1Δ* and *ssk1Δ* cells grown to exponential phase in YPD at 30°C. (C) Western blot analysis of whole cell extracts isolated from WT, *ssk1Δ,* and *ssk1Δ* +*SSK1* (JC1196) cells after treatment with 5mM H_2_O_2_, 0.1% SDS, 1M NaCl, 0.3M NaCl, or 1mM CdSO4 for the specified times. Western blots were probed with an anti-phospho-p38 antibody which recognises the phosphorylated form of Hog1 (Hog1-P). Total Hog1 levels were determined by stripping and reprobing the blot with an anti-Hog1 antibody that recognises phosphorylated and unphosphorylated Hog1. (D) Western blot analysis of whole cell extracts isolated from WT (JC747), *ssk1Δ* (JC1519)*, ssk1Δ trx1Δ* (JC1554), and *ssk1Δtrx1Δ+SSK1* (JC1555) cells after treatment with 5mM H_2_O_2_ for the specified times. Blots were processed as in (C).

Hog1 also functions to repress the yeast to hyphal switch in *C. albicans* (14) with *hog1Δ* cells forming hyphae under non filament-inducing conditions (44). Similarly, cells lacking *SSK1* also displayed a filamentous phenotype under non-inducing conditions (Fig. 1B). Collectively these data show that cells lacking the response regulator Ssk1 share many overlapping phenotypes with *hog1Δ* cells, with the significant exception of sensitivity to osmotic stress inducing agents.

To investigate whether Ssk1-dependent stress-sensitive phenotypes were linked to impaired activation of the Hog1 SAPK, stress-induced Hog1 phosphorylation was examined in wild-type and *ssk1Δ* cells. In line with previous reports, Hog1 phosphorylation in response to H_2_O_2_-induced oxidative stress was significantly impaired in *ssk1Δ* cells (32) and restored upon reintegration of *SSK1* (Fig. 1C). Hog1 phosphorylation in response to Cd^++^-imposed heavy metal stress and SDS-stress was also impaired in cells lacking Ssk1 (Fig. 1C).

Interestingly, in regard to osmotic stress, Ssk1 was required for Hog1 phosphorylation in response to low (0.3M) but not high (1M) levels of NaCl stress (Fig. 1C). This is consistent with the lack of stress-sensitivity of *ssk1Δ* cells in the presence of 1M NaCl (Fig. 1A), and previous work showing Ssk1 is dispensable for Hog1 activation following 1.5M NaCl stress (32). However, the observation that Ssk1 is required for Hog1 activation in response to low but not high levels of NaCl, indicates that concentration-dependent mechanisms relay osmotic stress signals to Hog1 in *C. albicans*. Taken together, the data presented here indicate that, with the exception of high levels of osmotic/salt stress, Ssk1 is a major regulator of the Hog1 SAPK in *C. albicans*.

Previous work from our group demonstrated that the thioredoxin redox protein Trx1 is important for oxidative stress-mediated activation of Hog1 (45). To explore whether this was acting in the same pathway as the Ssk1 response regulator, a double *ssk1Δtrx1Δ* mutant was made and oxidative-stress induced Hog1 activation examined. As illustrated in Fig. 1D, Hog1 activation in *ssk1Δtrx1Δ* cells was further diminished compared to the single *ssk1Δ* mutant, and reintegration of *SSK1* into *ssk1Δtrx1Δ* cells only partially rescued Hog1 activation. This indicates that Ssk1 and Trx1 work in independent pathways to regulate oxidative stress mediated activation of Hog1.

### Two component-dependent and -independent roles of Ssk1 in Hog1 regulation

Phosphorylation of Ssk1 via two-component signalling is mediated by the upstream phosphorelay protein Ypd1. Previously, we showed that in cells lacking Ypd1, where Ssk1 is trapped in the unphosphorylated form, Hog1 is hyper-activated even in the absence of stress. This hyper-activation was abolished in *ypd1Δssk1Δ* cells, confirming that Ypd1 acts solely through Ssk1 to activate Hog1 (30). If Ypd1-dependent regulation of Ssk1 phosphorylation is the primary mechanism underlying Ssk1-mediated regulation of Hog1, then no further stress-induced increases in Hog1 phosphorylation should be observed in *ypd1Δ* cells. To address this, we examined Hog1 phosphorylation in *ypd1Δ* cells in response to low NaCl and H_2_O_2_ stress (Fig. 2), which both require Ssk1 (Fig. 1C). As seen previously (30), high basal levels of Hog1 phosphorylation are seen in *ypd1Δ* cells in the absence of stress (Fig. 2, zero timepoints). However, following exposure to 0.3M NaCl, no further stress-induced increases in Hog1 phosphorylation were detected (Fig. 2A). This indicates that the role of Ssk1 in relaying a low osmotic stress signal to Hog1 is dependent on two-component signalling. In contrast, clear increases in stress induced Hog1 phosphorylation were observed in *ypd1Δ* cells in response to 5mM H_2_O_2_ (Fig. 2B). This suggests that Ssk1 relays oxidative stresses to Hog1 via a two-component independent mechanism.

**Figure 2.**
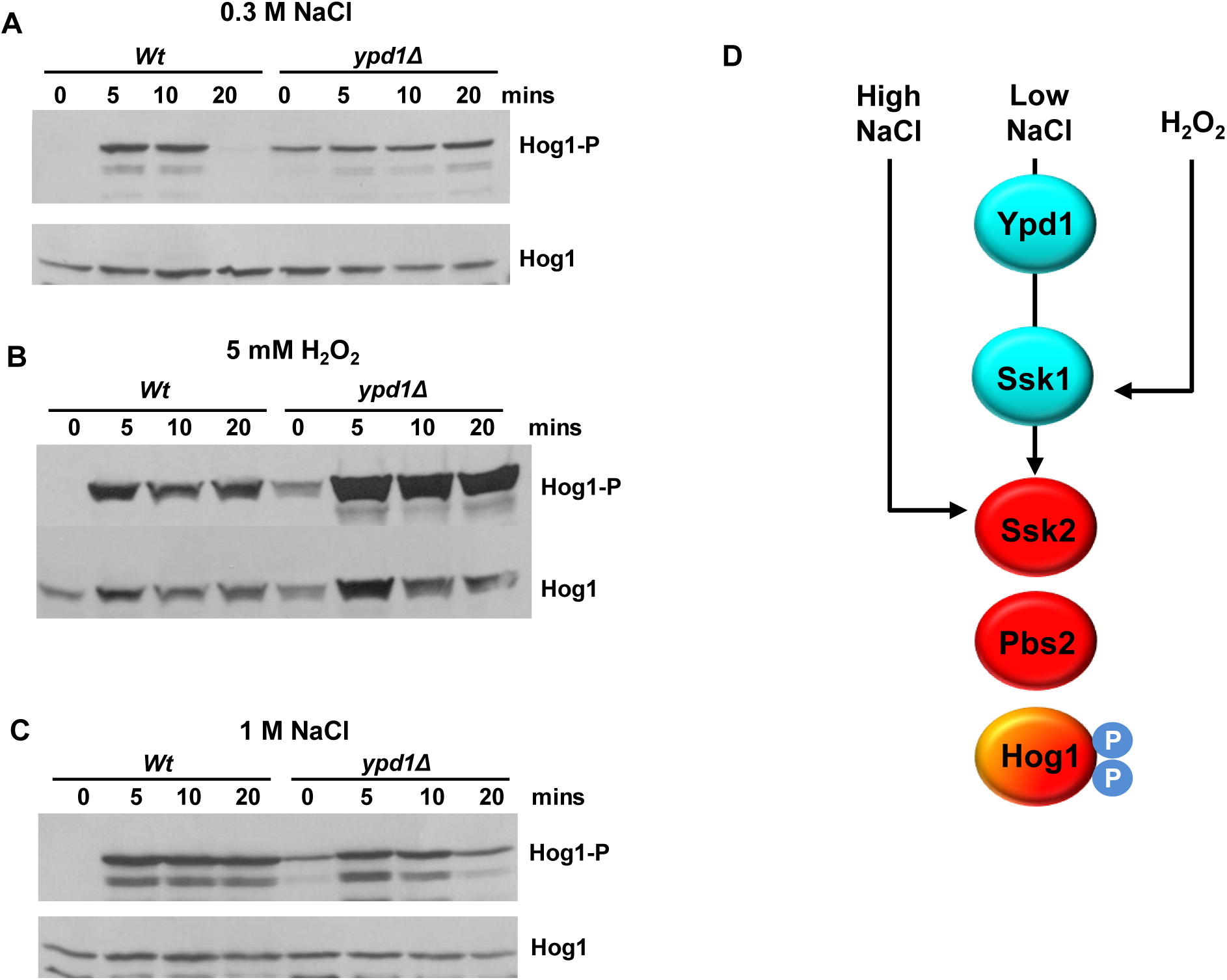
Stress conditional two-component dependent activation of Hog1. (A). Western blot analysis of whole cell extracts isolated from WT (JC26) and *ypd1Δ* (JC2001) cells after treatment with 0.3M NaCl (A), 1M NaCl (B), 5mM H_2_O_2_ (C) for the specified times. Western blots were probed with an anti-phospho-p38 antibody which recognises the phosphorylated form of Hog1 (Hog1-P). Total Hog1 levels were determined by stripping and reprobing the blot with an anti-Hog1 antibody that recognises phosphorylated and unphosphorylated Hog1. (D) Model summary. Activation of Hog1 in response to low NaCl requires Ypd1 and Ssk1, activation of Hog1 in response to H_2_O_2_ requires Ssk1 but not Ypd1, and activation of Hog1 in response to high NaCl requires neither Ypd1 nor Ssk1.

Hog1 phosphorylation was also examined in *ypd1Δ* cells in response to 1M NaCl (Fig. 2C). Consistent with Ssk1 being dispensable for Hog1 activation following 1M NaCl stress, stress-induced increases in Hog1 phosphorylation were observed in *ypd1Δ* cells. These data indicate three modes of Hog1 activation (summarised in Figure 2D): Ssk1 independent (1M NaCl); Ssk1 dependent and two-component dependent (0.3M NaCl); and Ssk1 dependent but two-component independent (H_2_O_2_). Thus, we set out to investigate the two-component independent mechanism of Hog1 regulation seen in response to H_2_O_2_ stress.

### Ssk1 functions as a scaffold for the interaction between Ssk2 and Pbs2

Activation of Hog1 in *C. albicans* is dependent on the upstream Pbs2 MAPKK which, in turn, is phosphorylated and activated by the upstream Ssk2 MAPKKK (25, 26). As Ssk1 regulates the Hog1 SAPK in response to oxidative stress via a mechanism independent of two-component signalling (Fig. 2), we hypothesised that this response regulator may also have a structural role. Thus, we explored whether Ssk1 functions as a scaffold protein, mediating interactions between the Pbs2 and Ssk2 upstream activating kinases. To facilitate this, wild-type and *ssk1Δ* mutant strains were constructed that express HA-epitope tagged Ssk2 and Myc-epitope tagged Pbs2. Pbs2 was immunoprecipitated using anti-myc antibody coupled agarose, and co-precipitation of Ssk2-HA was detected by western blotting. As illustrated in Figure 3A, Ssk2 does form a complex with Pbs2 in the absence of stress and, notably, this interaction is abolished in cells lacking the Ssk1 response regulator. This is consistent with the idea that Ssk1 functions as a scaffolding protein, promoting interactions between the Pbs2 and Ssk2 kinases in the Hog1 SAPK module. Next, we explored whether Pbs2-Ssk2 interactions were maintained following exposure to stresses that either require Ssk1 function for Hog1 activation (5 mM H_2_O_2_), or which activate Hog1 independently of Ssk1 (1 M NaCl). Strikingly, the interaction between Ssk2 and Pbs2 was maintained in an Ssk1-dependent manner following exposure to 5 mM H_2_O_2_, but completely abolished in response to 1M NaCl stress (Fig. 3A). We also investigated whether the interaction between Pbs2 and Ssk2 was disrupted following exposure to low levels of NaCl. In contrast to that seen with 1M NaCl, interaction between Pbs2 and Ssk2 was maintained following 0.3M NaCl stress, and again this was dependent on Ssk1 (Fig. 3B). Collectively, these data indicate that protein-protein interactions within the *C. albicans* Hog1 signalling module change in a stress-specific manner. The Ssk1-dependent interaction between Pbs1 and Ssk2 is maintained following exposure to H_2_O_2_ and 0.3M NaCl stresses and disrupted in response to 1M NaCl.

**Figure 3.**
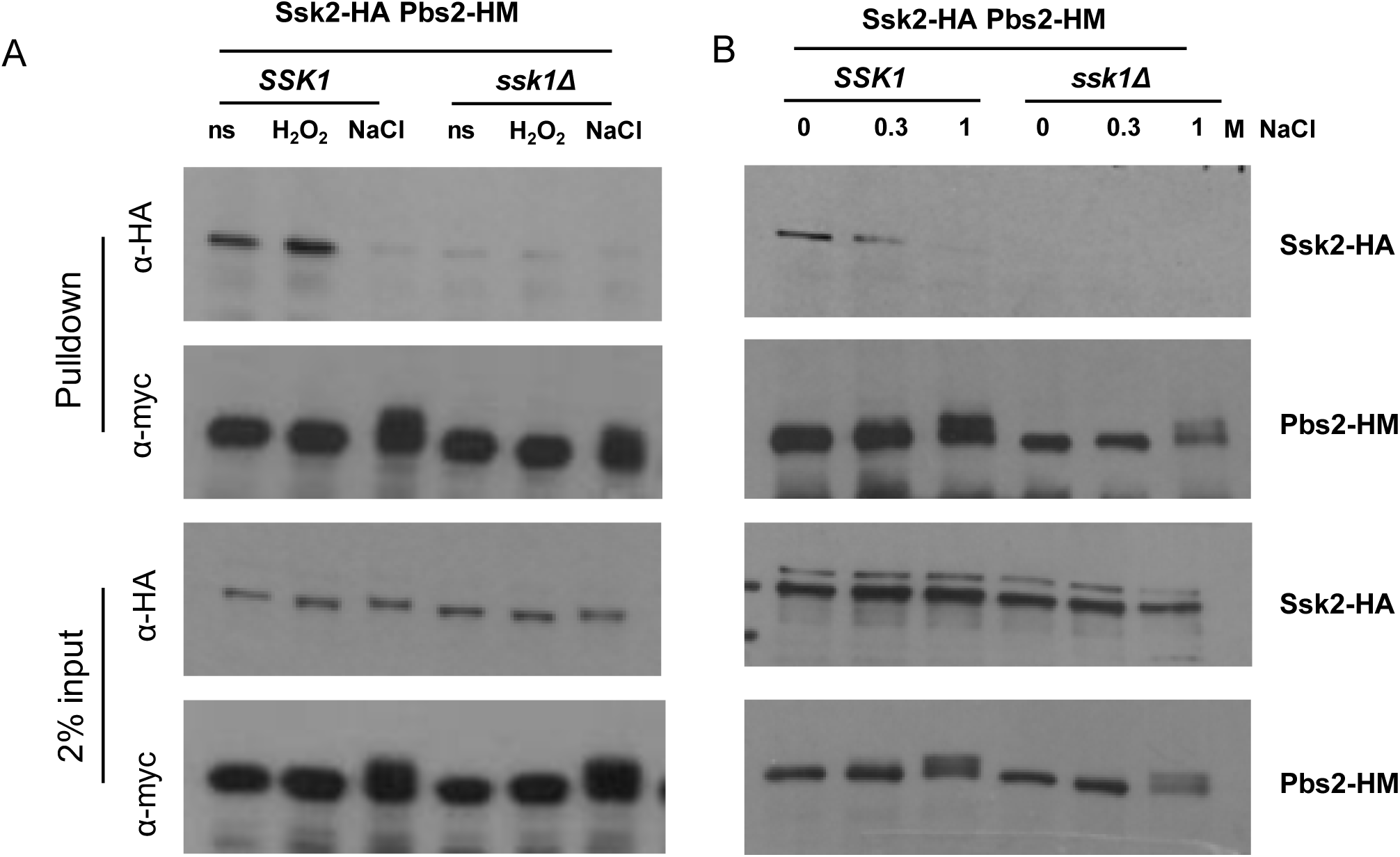
Scaffold function of Ssk1 and stress dependent changes in Hog1 pathway architecture. (A) Co-immunoprecipitation experiments to examine the association of Ssk2 and Pbs2. Extracts were prepared from *Wt* (JC2017) and *ssk1Δ* (JC2012) cells expressing Ssk2-HA and Pbs2-6His-myc (Pbs2-HM), before and after exposure to the stress conditions indicated. Pbs2-HM was precipitated using anti-myc agarose and co-immunoprecipitation of Ssk2-HA was analysed by Western blotting using an anti-HA antibody. Blots were stripped and reprobed with anti-myc to determine the levels of Pbs2-HM precipitated. 1% of protein input was also analysed by Western blotting using both anti-HA and anti-myc antibodies to determine amounts of Ssk2-HA and Pbs2-6His-myc in extracts used for co-immunoprecipitations.

### Phosphorylation of Pbs2 triggers its dissociation from Ssk2 in response to high NaCl

The above observations raised the question as to what drives Pbs2 dissociation from Ssk2 in response to high NaCl, but not H_2_O_2_ or low NaCl stresses. Pbs2 is activated via phosphorylation of conserved activating residues Ser355 and Thr359 (26), and stress-induced phosphorylation of Pbs2 is seen following high NaCl stress, as detected by the presence of slower migrating forms that are sensitive to phosphatase treatment (Fig 4A). Interestingly, similar mobility shifts of Pbs2 were not evident following exposure to H_2_O_2_ or low NaCl stresses suggesting that Pbs2 phosphorylation (and therefore activation) is less strong in response to these stresses compared to high NaCl stress. To test whether phosphorylation of Pbs2 drives its dissociation from Ssk2 following high NaCl stress, co-precipitation experiments were repeated in cells expressing Pbs2AA-myc, in which the activating phosphorylation sites (Ser355 and Thr359) are mutated to alanine (26). As illustrated in Fig. 4B, preventing Pbs2 phosphorylation strengthens the interaction of Ssk2 with Pbs2 under basal conditions and, importantly, inhibits the stress-induced dissociation of Pbs2 from Ssk2 in response to high NaCl stress. Taken together, these data indicate that phosphorylation of Pbs2 drives its dissociation from Ssk2 in response to high salt conditions.

**Figure 4.**
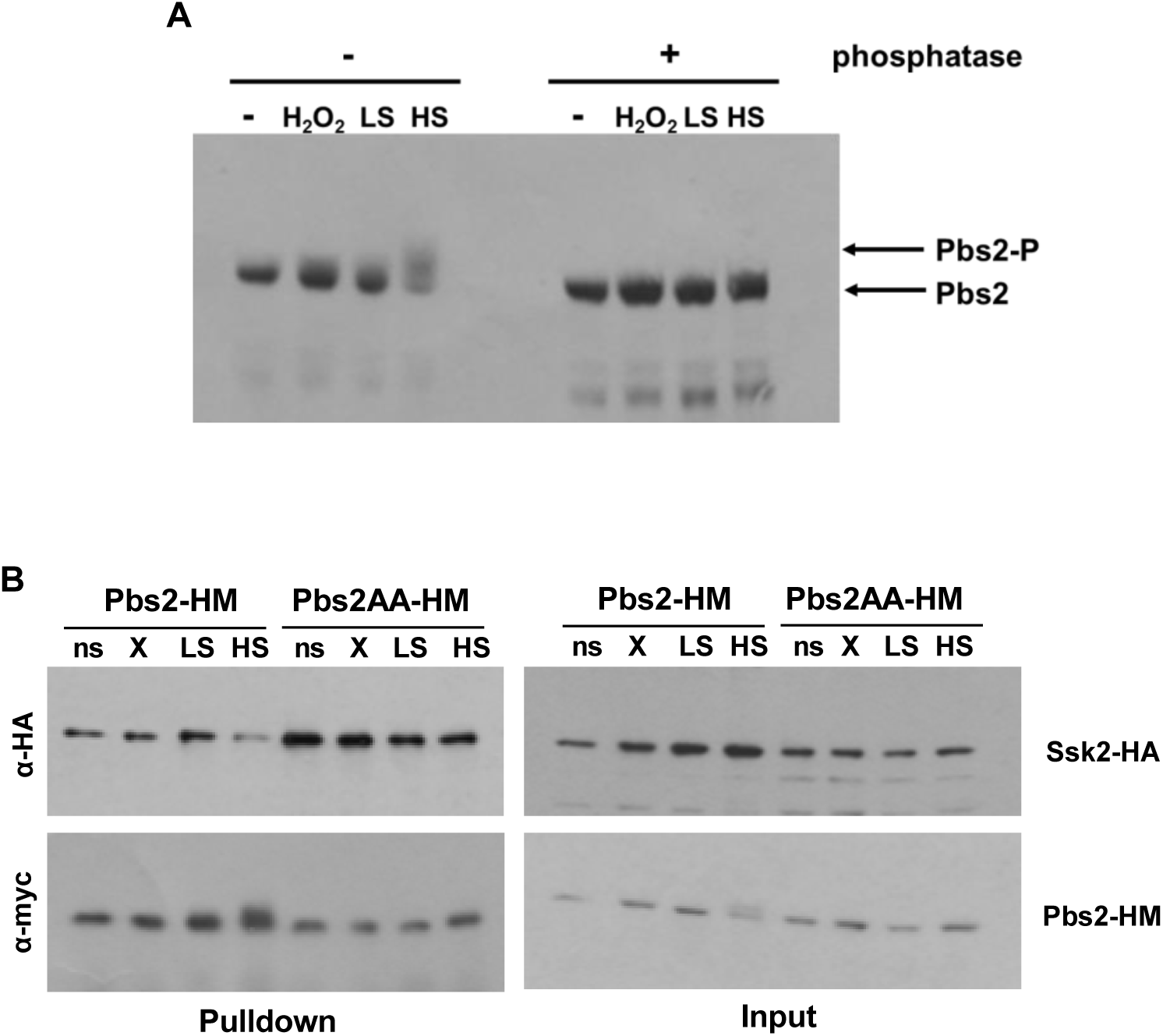
Phosphorylation of Pbs2 following high NaCl stress promotes its release from the Ssk2 MAPKKK. (A) Western blot analysis of whole cell extracts from *Wt* cells expressing Pbs2-HM (JC112) before (ns) and after stress for 10min with 5mM H_2_O_2_ (X), 0.3M NaCl (LS) or 1M NaCl (HS). Extracts were treated with or without λ phosphatase as indicated, and the blot was probed with an α-myc antibody. (B) Co-immunoprecipitation experiments to examine the association of Ssk2 with Pbs2 or Pbs2^AA^. Extracts were prepared from cells expressing Ssk2-HA and either Pbs2-HM (JC2121) or Pbs2^AA^-HM (JC2119), before (ns) or after exposure for 10 min to 5mM H_2_O_2_ (X), 0.3M NaCl (LS) or 1M NaCl (HS). Pbs2-HM or Pbs2^AA^-HM was precipitated using anti-myc antibody coupled agarose and co-immunoprecipitation of Ssk2-HA was analysed by Western blotting using an anti-HA antibody. Blots were stripped and reprobed with an anti-myc antibody to determine the levels of Pbs2-HM or Pbs2^AA^-HM precipitated. 1% of protein input was also analysed by Western blotting using both anti-HA and anti-myc antibodies to determine amounts of Ssk2-HA and Pbs2-HM/ Pbs2^AA^-HM in the extracts used for the co-immunoprecipitations.

### Hog1 activation in response to oxidative stress is mediated through inhibition of the protein tyrosine phosphatases Ptp2 and Ptp3

Hog1 is activated following exposure to either high NaCl or H_2_O_2_ stress (Fig. 1C). However, stress-induced activation of the Pbs2 upstream kinase is only detected with high NaCl stress. Therefore, what mechanism underlies oxidative stress mediated increases in Hog1 phosphorylation? In model yeasts, inactivation of downstream negative regulators has been implicated in Hog1 activation following certain stresses. For example, in *S. cerevisiae*, Hog1 activation in response to arsenite requires its metabolism to methyl arsenite which subsequently inhibits the protein tyrosine phosphatases (Ptp2 and Ptp3) that normally maintain Hog1 in an inactive state (46). In *S. pombe*, heat stress dissociates the protein tyrosine phosphatase Pyp1 from the Hog1 homologue, Sty1, which leads to strong Sty1 activation (47). Furthermore, of relevance here, oxidation and inactivation of Pyp1 in *S. pombe* following H_2_O_2_ stress contributes to Sty1 activation (48). In *C. albicans*, the Ptp2 and Ptp3 protein tyrosine phosphatases have been shown to function redundantly to dephosphorylate and inhibit Hog1 (27), thus we investigated their role in oxidative-stress mediated activation of this SAPK pathway.

Cells lacking *ptp2Δptp3Δ* were found to be more resistant to H_2_O_2_ stress, but not NaCl stress, linking the regulation of these phosphatases to oxidative stress signalling in *C. albicans* (Fig. 5A). Thus, we examined Hog1 phosphorylation in *C. albicans ptp2Δptp3Δ* cells following exposure to high NaCl or H_2_O_2_ stresses. If stress-induced inhibition of these phosphatases is a major driver of Hog1 activation, then any stress-induced increases in Hog1 activation should be diminished in *ptp2Δptp3Δ* cells. Consistent with the inhibitory action of Ptp2 and Ptp3, a high basal level of Hog1 phosphorylation was observed in cells lacking these phosphatases (Fig. 5B). In response to both osmotic and oxidative stress, the fold induction of Hog1 activation was reduced in *ptp2Δptp3Δ* cells compared to wild-type cells (Fig. 5B). However, this reduction was greater following oxidative stress where only a 2.5-fold increase in Hog1 phosphorylation was seen in *ptp2Δptp3Δ* cells after 10 min H_2_O_2_ stress, compared to a 9-fold increase after 10 min NaCl stress. This supports the hypothesis that H_2_O_2_-mediated inhibition of these phosphatases plays a major role in Hog1 activation following oxidative stress, whereas phosphorylation of Hog1 in response to high salt is instead mediated via activation of the upstream Pbs2 kinase.

**Figure 5.**
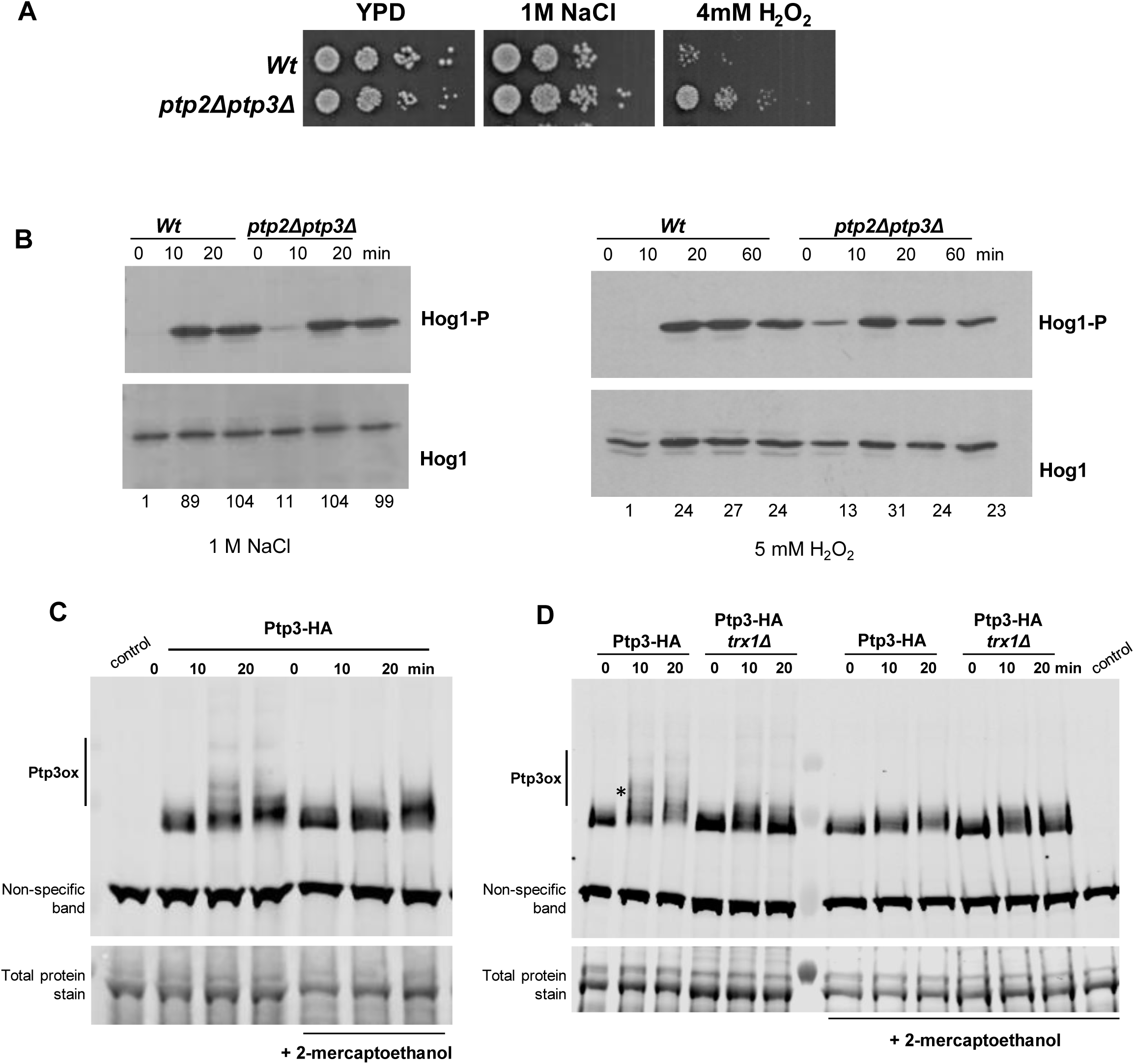
Role of the protein tyrosine phosphatases Ptp2 and Ptp3 in oxidative stress responses and Hog1 regulation. (A) Exponentially growing *Wt* (SN250) or *ptp2Δptp3Δ* (JC2291) cells were serially diluted 10-fold and spotted onto YPD plates containing 1M NaCl or 4mM H_2_O_2_. Plates were incubated at 30°C for 48h. (B) Western blot analysis of Hog1 phosphorylation in whole cell extracts isolated from *Wt* or *ptp2Δptp3Δ* cells after treatment with 1M NaCl or 5mM H_2_O_2_ for the specified times. Western blots were probed with an anti-phospho-p38 antibody which recognises the phosphorylated form of Hog1 (Hog1-P). Total Hog1 levels were determined by stripping and reprobing the blot with an anti-Hog1 antibody that recognises phosphorylated and unphosphorylated Hog1. Blots were quantified using Image J (Version 1.54) and levels of Hog1 phosphorylation in each strain after stress are expressed relative to the level of Hog1 phosphorylation in *Wt* cells before stress. (C) Western blot analysis of Ptp3 oxidation state in wild-type (JC2231) cells expressing HA-tagged Ptp3 (Ptp3-HA) before and following treatment with 5 mM H_2_O_2_ for the specified times. Samples were ran under both non-reducing and reducing (+ 2-mercaptoethanol) conditions to determine oxidised forms of Ptp3. A loading control (total protein stain) is also shown. (D) Western blot analysis of Ptp3 oxidation state in wild-type (JC2231) and *trx1Δ* (JC2626) cells expressing HA-tagged Ptp3 (Ptp3-HA) before and following treatment with 5 mM H_2_O_2_ for the specified times. Samples were analysed as in (C). The asterisk highlights the oxidised form of Ptp3 that is dependent on Trx1.

To explore whether oxidation could underlie inactivation of the protein tyrosine phosphatases, akin to that seen in *S. pombe*, we examined whether Ptp3 was oxidised following H_2_O_2_ stress. Exposure of cells expressing HA-tagged Ptp3 to H_2_O_2_ revealed the presence of several slower-migrating forms of Ptp3 (Fig. 5C). To determine if the slower-migrating forms were due to oxidation, samples were reduced with 2-mercaptoethanol prior to SDS-PAGE. Although some 2-mercaptothanol-resistant slower migrating forms remain, other slower-migrating forms were absent following 2-mercaptoethanol treatment thus identifying these as oxidized forms of Ptp3 (Figure 5C). The Trx1 thioredoxin oxidoreductase is important for protein-tyrosine phosphatase oxidation in fission yeast (48) and, notably, Trx1 is also essential for H_2_O_2_-mediated activation of Hog1 in *C. albicans* (45), in an Ssk1-independent mechanism (Fig. 1D). Thus, we examined Ptp3 oxidation in *C. albicans trx1Δ* cells. In contrast to that seen in wild-type cells, specific oxidised forms of Ptp3 were absent in cells lacking Trx1 showing that this oxidative stress-induced modification is dependent on Trx1 (Fig. 5D). Taken together these data suggest that H_2_O_2_-mediated oxidation of the protein tyrosine phosphatases lowers their activity against Hog1 resulting in increases in Hog1 phosphorylation. Furthermore, the observation that Trx1 is contributes to the oxidation and inactivation of these phosphatases is consistent with the previously described role of this oxidoreductase in H_2_O_2_-induced-Hog1 activation (45).

## Discussion

In this study we reveal stress contingent mechanisms of Hog1 regulation in *C. albicans* (Fig. 6). Central to this is our finding that the Ssk1 response regulator functions as a scaffolding protein that promotes interactions between the Ssk2 and Pbs2 kinases that lie upstream of Hog1 within this MAPK pathway (Fig. 3). Following osmotic stress, the Pbs2 MAPKK becomes robustly phosphorylated and this drives dissociation of Pbs2 from Ssk2. However, following oxidative stress, limited phosphorylation of Pbs2 is observed and the Ssk1-mediated Ssk2 and Pbs2 interaction is maintained (Fig. 4). Here, we present evidence that oxidative stress-mediated increases in Hog1 activation are due to the oxidation and inhibition of the phosphatases Ptp2 and Ptp3, that negatively regulate Hog1 (Fig. 5). We propose that the Ssk1-scaffolding function is required to maintain a basal flux through the Hog1 SAPK pathway which is important for responses to stresses such as oxidative stress that do not stimulate robust activation of the upstream Ssk1 and Pbs2 kinases. As Ssk1 is required for Hog1 activation in response to many diverse stresses in *C. albicans*, this alternative mode of stress-mediated increases in Hog1 phosphorylation may be applicable to other stresses. Basal levels of Hog1 phosphorylation are also important to repress filamentation under non-hyphae inducing conditions, and accordingly *ssk1Δ* cells exhibit the same morphological defects as cells lacking Hog1 (Fig. 1) or expressing a non-phosphorylatable Hog1^AF^ mutant (19).

**Fig 6.**
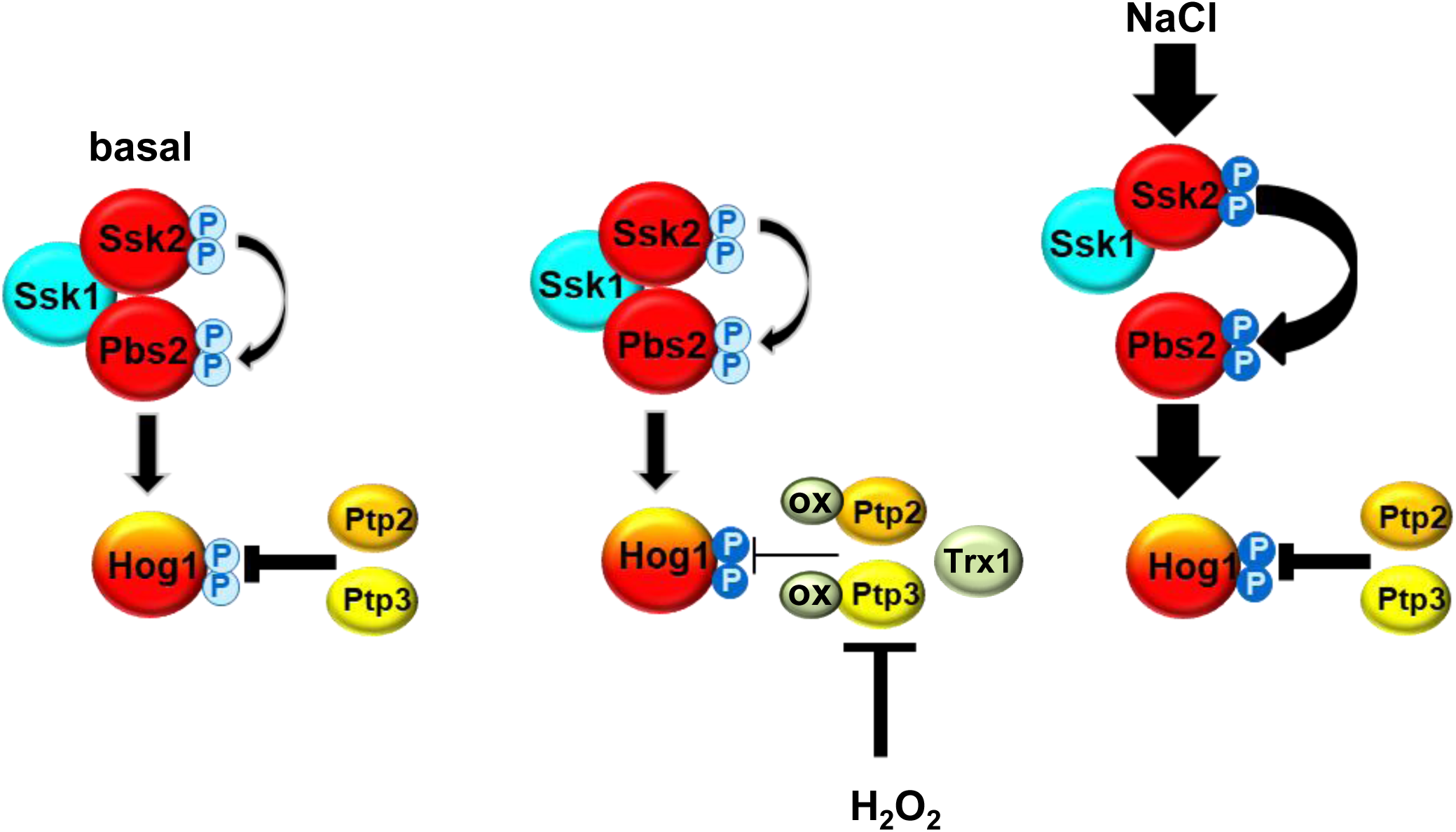
Model depicting stress contingent changes in Hog1 pathway architecture and regulation in *C. albicans*. Under non stressed conditions a basal level of Hog1 phosphorylation is dynamically maintained by the opposing actions of the positive regulators upstream and negative regulators downstream. Specifically, this involves the scaffolding function of Ssk1 which facilitates a close association between Ssk2 and Pbs2 kinases promoting flux through the pathway, and the opposing the action of the protein tyrosine phosphatases Ptp2 and Ptp3. Following oxidative stress, the Ssk1 scaffold is needed to promote Ssk2 and Pbs2 interaction and basal flux through the pathway. However, rather than activation of upstream kinases, stress-induced increases in Hog1 activation involve the Trx1-dependent oxidation and inactivation of the protein tyrosine phosphatases that negatively regulate Hog1. Finally, in response to high NaCl the Pbs2 kinase becomes robustly phosphorylated driving Hog1 activation. Here the Ssk1 scaffold is not needed; indeed, phosphorylation of Pbs2 drives its dissociation from Ssk2.

The response regulator Ssk1 is conserved across fungal species (23). In *S. cerevisiae*, deletion of Ssk1 does not impact on Hog1 regulation as the Sho1 osmotic stress signalling pathway functions in parallel with the Sln1-Ypd1-Ssk1 two component pathway (49). However, in cells lacking the Sho1 signalling branch, osmotic stress activation of Hog1 is mediated via inhibition of the Sln1 histidine kinase, which results in unphosphorylated Ssk1, a potent activator of the Ssk2/22 MAPKKKs (50). In *C. albicans ypd1Δ* cells, where Ssk1 is locked in the unphosphorylated form, no further Hog1 activation is observed in response to low levels of cationic stress (Fig. 2). This is consistent with the Sln1 histidine kinase acting as an osmosensor that regulates Hog1 activation through the phosphorylation status of the Ssk1 response regulator. It is probable that the *C. albicans* Sln1 histidine kinase will also respond to high levels of osmotic stress, but this is masked by a second, as yet uncharacterised, osmosensing pathway that functions independently of Ssk1 (see below).

In *S. pombe*, the Ssk1 homologue (Mcs4) also regulates activation of the Sty1 SAPK by two-component dependent and two-component independent mechanisms. Two-component mediated phosphorylation of Mcs4 is important for the relay of oxidative stress signals to Sty1 (51), whereas Mcs4 regulates osmotic and heat stress activation of Sty1 in a two-component independent mechanism (51, 52). Here, Mcs4 functions to promote heteromer formation of the two non-redundant MAPKKKs Wak1 and Win1, which is essential for the relay of stress signals to Sty1 (53). In *C. albicans*, Hog1 is regulated by a single MAPKKK, Ssk2, and, in contrast to that seen in *S. pombe*, osmotic stress-mediated activation of Hog1 is Ssk1-independent, but Ssk2 dependent (26, 32). Thus, there appear to be differences between *C. albicans* and *S. pombe* with respect to the mechanisms underlying the two-component independent roles of the Ssk1/Mcs4 response regulators in SAPK pathway regulation. Nonetheless, homologues of the Ssk1 response regulator also play global roles in Hog1 activation in other pathogenic fungi including *Cryptococcus neoformans* (54), *Candida auris* (55) and the plant pathogens *Alternaria alternata* (56) and *Cochliobolus heterostrophus* (57), with *ssk1Δ* cells largely phenocopying *hog1Δ* cells in these organisms. Thus, a scaffolding function of Ssk1, that promotes key interactions within the three-tiered SAPK module, may contribute to the global role of Ssk1 in SAPK regulation in other fungi.

Regarding osmotic stress signalling to Hog1 in *C. albicans,* two questions arise from this study; what is the Ssk1-independent mechanism of osmo-signalling to Hog1, and why does the Ssk1-mediated Pbs2-Ssk2 scaffold disassemble following osmotic stress? In *S. cerevisiae* the binding of unphosphorylated Ssk1 to the autoinhibitory domain of Ssk2 relieves the inhibition resulting in the autophosphoryation and activation of the MAPKKK (31). As Pbs2 is robustly phosphorylated following high levels of osmotic stress in *C. albicans,* a signalling mechanism must drive the Ssk1-independent activation of the upstream Ssk2 MAPKKK. Interestingly, in *S. cerevisiae* cells engineered to remove both upstream osmo-signalling pathways, Ssk2-dependent activation of Hog1 is seen at high but not low levels of osmotic stress. The domain responsible for the Ssk1-independent activation of Ssk2 was mapped between amino acid residues 177∼240, which is distinct from the Ssk1 binding domain (residues 294–413), leading to the suggestion that this region forms a binding site for an unknown regulator which in turn activates Ssk2 (58). However, in *C. albicans,* osmotic stress-induced increases in Hog1 phosphorylation are observed in cells expressing a truncated version of Ssk2 lacking the entire N-terminal non-catalytic domain (19). Thus, the unknown osmotic stress regulator(s) of the Ssk2 MAPKKK in *C. albicans* may target the kinase domain directly. Regarding the disassembly of the Ssk1-Ssk2-Pbs2 complex following osmotic stress, one possibility is that this facilitates the interaction of Pbs2 with its substrate Hog1. However, we could find no evidence of increased Pbs2-Hog1 interaction in response to high salt stress. In *S. cerevisiae*, in response to high levels of osmotic stress the actin cytoskeleton rapidly disassembles thus resulting in a stalling of the cell cycle (59). Reassembly of the actin cytoskeleton is expediated by Ssk2 which localises to the bud neck in response to osmotic stress and binds to actin (60, 61). Such a response is conserved in mammalian cells as the MAPKKK on the p38 and JNK SAPK pathways (MEKK4), is similarly required for actin reassembly following osmotic stress (62). Future work could explore whether disassembly of the Ssk2-Pbs2 complex in *C. albicans*, specifically in response to high osmotic stress, facilitates Hog1 independent functions of Ssk2 such as reassembly of the actin cytoskeleton.

A further question raised by this study centres around the importance of stress-mediated increases in Pbs2 phosphorylation in Hog1 activation. Ssk2 phosphorylates Pbs2 on Ser355 and Thr359, and mutation of these residues blocks Hog1 activation in response to all stresses tested (19). However, in contrast to the dogma that stress-mediated Hog1 phosphorylation is via activation of the upstream kinases, in this study we failed to detect significant increases in Pbs2 phosphorylation following oxidative stress. Under such conditions, we suggest that the Ssk1-mediated tight association of Ssk2 and Pbs2 is important to maintain a basal level of signalling to Hog1, with stress-induced increases in Hog1 activation occurring independently of Pbs2/Ssk2. This mode of stress-mediated increases in Hog1 phosphorylation may be applicable to other stresses as Ssk1 is required for Hog1 activation in response to many diverse stresses in *C. albicans* (Fig.1). Focussing on oxidative stress signalling, whilst notable increases in Pbs2 phosphorylation were not observed, we show that inactivation of the negative regulators Ptp2 and Ptp3 drives increases in oxidative stress mediated Hog1 phosphorylation. This is consistent with work in other organisms showing that protein tyrosine kinases are susceptible to oxidative-stress mediated inhibition through oxidation of the active site cysteine residue (63). Furthermore, a recent study in the fission yeast *S. pombe*, revealed that oxidation of the Pyp1 protein tyrosine phosphatase to slower migrating disulfide-bonded complexes, contributes to increases in Sty1 phosphorylation (48). Excitingly, we find that the *C. albicans* Ptp3 phosphatase is oxidised to slower migrating disulfide-bonded complexes following oxidative stress. Moreover, as reported in fission yeast (48) we find that the thioredoxin protein, Trx1, is important for Ptp3 oxidation (Fig. 5C) which likely underlies the importance of Trx1 in mediating H_2_O_2_-induced phosphorylation of Hog1 in *C. albicans* (45). Collectively, these data support a model of Hog1 activation in response to oxidative stress that is dependent on the scaffolding function of Ssk1 combined with Trx1-mediated oxidation and inactivation of the Ptp2/3 negative regulators. Our data that Trx1 and Ssk1 regulate Hog1 independently (Fig. 1D) further supports this model.

Hog1-related SAPKs are important virulence determinants in many fungal pathogens. However, given the high degree of functional and structural homology between fungal and host SAPKs (22), it may be challenging to design antifungals that are specific for fungal Hog1. Instead, Ssk1 is a fungal-specific SAPK regulator that has been shown to be an important virulence factor of many human and plant fungal pathogens. Therefore, understanding how this response regulator regulates SAPKs may inform the development of Ssk1-targetting antifungals.

## Acknowledgements

We thank Professor Haoping Liu for the kind gift of the *ptpΔ2ptp3Δ* deletion strain. JQ was funded by the BBSRC [BB/K016393/1, BB/P020119/1], and Wellcome [215599/Z/19/Z].

AJPB was funded by the BBSRC [BB/K017365/1], the MRC [MR/M026663/1, MR/M026663/2] and Wellcome [224323/Z/21/Z] and was supported by the Medical Research Council Centre for Medical Mycology [MR/N006364/1, MR/N006364/2].

## Notes

### Competing Interest Statement

The authors have declared no competing interest.

## References

1. Fisher MC, Denning DW. The WHO fungal priority pathogens list as a game-changer. Nat Rev Microbiol. 2023;21(4):211–2.

2. Limon JJ, Skalski JH, Underhill DM. Commensal Fungi in Health and Disease. Cell Host Microbe. 2017;22(2):156–65.

3. Wall G, Lopez-Ribot JL. Current Antimycotics, New Prospects, and Future Approaches to Antifungal Therapy. Antibiotics (Basel). 2020;9(8).

4. Lee Y, Puumala E, Robbins N, Cowen LE. Antifungal Drug Resistance: Molecular Mechanisms in Candida albicans and Beyond. Chem Rev. 2021;121(6):3390–411.

5. Koehler P, Stecher M, Cornely OA, Koehler D, Vehreschild M, Bohlius J, et al. Morbidity and mortality of candidaemia in Europe: an epidemiologic meta-analysis. Clin Microbiol Infect. 2019;25(10):1200–12.

6. Tsay SV, Mu Y, Williams S, Epson E, Nadle J, Bamberg WM, et al. Burden of Candidemia in the United States, 2017. Clin Infect Dis. 2020;71(9):e449–e53.

7. Brown AJP, Cowen LE, di Pietro A, Quinn J. Stress Adaptation. Microbiology spectrum. 2017;5(4).

8. Alves R, Barata-Antunes C, Casal M, Brown AJP, Van Dijck P, Paiva S. Adapting to survive: How Candida overcomes host-imposed constraints during human colonization. PLoS pathogens. 2020;16(5):e1008478.

9. Amulic B, Cazalet C, Hayes GL, Metzler KD, Zychlinsky A. Neutrophil function: from mechanisms to disease. Annu Rev Immunol. 2012;30:459–89.

10. Bensen ES, Martin SJ, Li M, Berman J, Davis DA. Transcriptional profiling in Candida albicans reveals new adaptive responses to extracellular pH and functions for Rim101p. Molecular microbiology. 2004;54(5):1335–51.

11. Ohno A, Müller E, Fraek ML, Thurau K, Beck F. Solute composition and heat shock proteins in rat renal medulla. Pflugers Arch. 1997;434(1):117–22.

12. Alselami A, Drummond RA. How metals fuel fungal virulence, yet promote anti-fungal immunity. Dis Model Mech. 2023;16(10).

13. Day AM, Quinn J. Stress-Activated Protein Kinases in Human Fungal Pathogens. Front Cell Infect Microbiol. 2019;9:261.

14. Alonso-Monge R, Navarro-Garcia F, Molero G, Diez-Orejas R, Gustin M, Pla J, et al. Role of the mitogen-activated protein kinase Hog1p in morphogenesis and virulence of Candida albicans. Journal of bacteriology. 1999;181(10):3058–68.

15. Liang SH, Cheng JH, Deng FS, Tsai PA, Lin CH. A novel function for Hog1 stress-activated protein kinase in controlling white-opaque switching and mating in Candida albicans. Eukaryotic cell. 2014;13(12):1557–66.

16. Alonso-Monge R, Carvaihlo S, Nombela C, Rial E, Pla J. The Hog1 MAP kinase controls respiratory metabolism in the fungal pathogen Candida albicans. Microbiology (Reading, England). 2009;155(Pt 2):413–23.

17. O’Meara TR, Duah K, Guo CX, Maxson ME, Gaudet RG, Koselny K, et al. High-Throughput Screening Identifies Genes Required for Candida albicans Induction of Macrophage Pyroptosis. mBio. 2018;9(4).

18. Sellam A, Chaillot J, Mallick J, Tebbji F, Richard Albert J, Cook MA, et al. The p38/HOG stress-activated protein kinase network couples growth to division in Candida albicans. PLoS genetics. 2019;15(3):e1008052.

19. Cheetham J, MacCallum DM, Doris KS, da Silva Dantas A, Scorfield S, Odds F, et al. MAPKKK-independent regulation of the Hog1 stress-activated protein kinase in Candida albicans. The Journal of biological chemistry. 2011;286(49):42002–16.

20. Prieto D, Roman E, Correia I, Pla J. The HOG pathway is critical for the colonization of the mouse gastrointestinal tract by Candida albicans. PloS one. 2014;9(1):e87128.

21. Arana DM, Alonso-Monge R, Du C, Calderone R, Pla J. Differential susceptibility of mitogen-activated protein kinase pathway mutants to oxidative-mediated killing by phagocytes in the fungal pathogen Candida albicans. Cellular microbiology. 2007;9(7):1647–59.

22. Herrero-de-Dios C, Day AM, Tillmann AT, Kastora SL, Stead D, Salgado PS, et al. Redox Regulation, Rather than Stress-Induced Phosphorylation, of a Hog1 Mitogen-Activated Protein Kinase Modulates Its Nitrosative-Stress-Specific Outputs. mBio. 2018;9(2).

23. Nikolaou E, Agrafioti I, Stumpf M, Quinn J, Stansfield I, Brown AJ. Phylogenetic diversity of stress signalling pathways in fungi. BMC evolutionary biology. 2009;9:44.

24. Zarubin T, Han J. Activation and signaling of the p38 MAP kinase pathway. Cell Res. 2005;15(1):11–8.

25. Arana DM, Nombela C, Alonso-Monge R, Pla J. The Pbs2 MAP kinase kinase is essential for the oxidative-stress response in the fungal pathogen Candida albicans. Microbiology (Reading, England). 2005;151(Pt 4):1033–49.

26. Cheetham J, Smith DA, da Silva Dantas A, Doris KS, Patterson MJ, Bruce CR, et al. A single MAPKKK regulates the Hog1 MAPK pathway in the pathogenic fungus Candida albicans. Molecular biology of the cell. 2007;18(11):4603–14.

27. Su C, Lu Y, Liu H. Reduced TOR signaling sustains hyphal development in Candida albicans by lowering Hog1 basal activity. Molecular biology of the cell. 2013;24(3):385–97.

28. Shor E, Chauhan N. A case for two-component signaling systems as antifungal drug targets. PLoS pathogens. 2015;11(2):e1004632.

29. Liao B, Ye X, Chen X, Zhou Y, Cheng L, Zhou X, et al. The two-component signal transduction system and its regulation in Candida albicans. Virulence. 2021;12(1):1884–99.

30. Day AM, Smith DA, Ikeh MA, Haider M, Herrero-de-Dios CM, Brown AJ, et al. Blocking two-component signalling enhances Candida albicans virulence and reveals adaptive mechanisms that counteract sustained SAPK activation. PLoS pathogens. 2017;13(1):e1006131.

31. Posas F, Saito H. Activation of the yeast SSK2 MAP kinase kinase kinase by the SSK1 two-component response regulator. The EMBO journal. 1998;17(5):1385–94.

32. Chauhan N, Inglis D, Roman E, Pla J, Li D, Calera JA, et al. Candida albicans response regulator gene SSK1 regulates a subset of genes whose functions are associated with cell wall biosynthesis and adaptation to oxidative stress. Eukaryotic cell. 2003;2(5):1018–24.

33. Roman E, Nombela C, Pla J. The Sho1 adaptor protein links oxidative stress to morphogenesis and cell wall biosynthesis in the fungal pathogen Candida albicans. Molecular and cellular biology. 2005;25(23):10611–27.

34. Menon V, Li D, Chauhan N, Rajnarayanan R, Dubrovska A, West AH, et al. Functional studies of the Ssk1p response regulator protein of Candida albicans as determined by phenotypic analysis of receiver domain point mutants. Molecular microbiology. 2006;62(4):997–1013.

35. Calera JA, Zhao XJ, Calderone R. Defective hyphal development and avirulence caused by a deletion of the SSK1 response regulator gene in Candida albicans. Infection and immunity. 2000;68(2):518–25.

36. Sherman F. Getting started with yeast. Methods in enzymology. 1991;194:3–21.

37. Bruce CR, Smith DA, Rodgers D, da Silva Dantas A, MacCallum DM, Morgan BA, et al. Identification of a novel response regulator, Crr1, that is required for hydrogen peroxide resistance in Candida albicans. PloS one. 2011;6(12):e27979.

38. Murad AM, Lee PR, Broadbent ID, Barelle CJ, Brown AJ. CIp10, an efficient and convenient integrating vector for Candida albicans. Yeast. 2000;16(4):325–7.

39. Noble SM, Johnson AD. Strains and strategies for large-scale gene deletion studies of the diploid human fungal pathogen Candida albicans. Eukaryotic cell. 2005;4(2):298–309.

40. Dennison PM, Ramsdale M, Manson CL, Brown AJ. Gene disruption in Candida albicans using a synthetic, codon-optimised Cre-loxP system. Fungal genetics and biology : FG & B. 2005;42(9):737–48.

41. Fonzi WA, Irwin MY. Isogenic strain construction and gene mapping in Candida albicans. Genetics. 1993;134(3):717–28.

42. Lavoie H, Sellam A, Askew C, Nantel A, Whiteway M. A toolbox for epitope-tagging and genome-wide location analysis in Candida albicans. BMC Genomics. 2008;9:578.

43. Smith DA, Nicholls S, Morgan BA, Brown AJ, Quinn J. A conserved stress-activated protein kinase regulates a core stress response in the human pathogen Candida albicans. Molecular biology of the cell. 2004;15(9):4179–90.

44. Enjalbert B, Smith DA, Cornell MJ, Alam I, Nicholls S, Brown AJ, et al. Role of the Hog1 stress-activated protein kinase in the global transcriptional response to stress in the fungal pathogen Candida albicans. Molecular biology of the cell. 2006;17(2):1018–32.

45. da Silva Dantas A, Patterson MJ, Smith DA, Maccallum DM, Erwig LP, Morgan BA, et al. Thioredoxin regulates multiple hydrogen peroxide-induced signaling pathways in Candida albicans. Molecular and cellular biology. 2010;30(19):4550–63.

46. Lee J, Levin DE. Intracellular mechanism by which arsenite activates the yeast stress MAPK Hog1. Molecular biology of the cell. 2018;29(15):1904–15.

47. Nguyen AN, Shiozaki K. Heat-shock-induced activation of stress MAP kinase is regulated by threonine- and tyrosine-specific phosphatases. Genes & development. 1999;13(13):1653–63.

48. Cao M, Day AM, Galler M, Latimer HR, Byrne DP, Foy TW, et al. A peroxiredoxin-P38 MAPK scaffold increases MAPK activity by MAP3K-independent mechanisms. Molecular cell. 2023;83(17):3140–54.e7.

49. Maeda T, Takekawa M, Saito H. Activation of yeast PBS2 MAPKK by MAPKKKs or by binding of an SH3-containing osmosensor. Science (New York, NY). 1995;269(5223):554-8.

50. Posas F, Wurgler-Murphy SM, Maeda T, Witten EA, Thai TC, Saito H. Yeast HOG1 MAP kinase cascade is regulated by a multistep phosphorelay mechanism in the SLN1-YPD1-SSK1 “two-component” osmosensor. Cell. 1996;86(6):865–75.

51. Buck V, Quinn J, Soto Pino T, Martin H, Saldanha J, Makino K, et al. Peroxide sensors for the fission yeast stress-activated mitogen-activated protein kinase pathway. Molecular biology of the cell. 2001;12(2):407–19.

52. Shieh JC, Wilkinson MG, Buck V, Morgan BA, Makino K, Millar JB. The Mcs4 response regulator coordinately controls the stress-activated Wak1-Wis1-Sty1 MAP kinase pathway and fission yeast cell cycle. Genes & development. 1997;11(8):1008–22.

53. Morigasaki S, Ikner A, Tatebe H, Shiozaki K. Response regulator-mediated MAPKKK heteromer promotes stress signaling to the Spc1 MAPK in fission yeast. Molecular biology of the cell. 2013;24(7):1083–92.

54. Bahn YS, Kojima K, Cox GM, Heitman J. A unique fungal two-component system regulates stress responses, drug sensitivity, sexual development, and virulence of Cryptococcus neoformans. Molecular biology of the cell. 2006;17(7):3122–35.

55. Shivarathri R, Jenull S, Stoiber A, Chauhan M, Mazumdar R, Singh A, et al. The Two-Component Response Regulator Ssk1 and the Mitogen-Activated Protein Kinase Hog1 Control Antifungal Drug Resistance and Cell Wall Architecture of Candida auris. mSphere. 2020;5(5).

56. Yu PL, Chen LH, Chung KR. How the Pathogenic Fungus Alternaria alternata Copes with Stress via the Response Regulators SSK1 and SHO1. PloS one. 2016;11(2):e0149153.

57. Oide S, Liu J, Yun SH, Wu D, Michev A, Choi MY, et al. Histidine kinase two-component response regulator proteins regulate reproductive development, virulence, and stress responses of the fungal cereal pathogens Cochliobolus heterostrophus and Gibberella zeae. Eukaryotic cell. 2010;9(12):1867–80.

58. Zhi H, Tang L, Xia Y, Zhang J. Ssk1p-independent activation of Ssk2p plays an important role in the osmotic stress response in Saccharomyces cerevisiae: alternative activation of Ssk2p in osmotic stress. PloS one. 2013;8(2):e54867.

59. Chowdhury S, Smith KW, Gustin MC. Osmotic stress and the yeast cytoskeleton: phenotype-specific suppression of an actin mutation. J Cell Biol. 1992;118(3):561–71.

60. Yuzyuk T, Amberg DC. Actin recovery and bud emergence in osmotically stressed cells requires the conserved actin interacting mitogen-activated protein kinase kinase kinase Ssk2p/MTK1 and the scaffold protein Spa2p. Molecular biology of the cell. 2003;14(7):3013–26.

61. Yuzyuk T, Foehr M, Amberg DC. The MEK kinase Ssk2p promotes actin cytoskeleton recovery after osmotic stress. Molecular biology of the cell. 2002;13(8):2869–80.

62. Bettinger BT, Amberg DC. The MEK kinases MEKK4/Ssk2p facilitate complexity in the stress signaling responses of diverse systems. J Cell Biochem. 2007;101(1):34–43.

63. Tonks NK. Redox redux: revisiting PTPs and the control of cell signaling. Cell. 2005;121(5):667–70.

64. Negredo A, Monteoliva L, Gil C, Pla J, Nombela C. Cloning, analysis and one-step disruption of the ARG5,6 gene of Candida albicans. Microbiology (Reading). 1997;143 ( Pt 2):297–302.

